# The retrotransposon *R2* maintains *Drosophila* ribosomal DNA repeats

**DOI:** 10.1101/2021.07.12.451825

**Authors:** Jonathan O. Nelson, Alyssa Slicko, Yukiko M. Yamashita

## Abstract

Ribosomal DNA (rDNA) loci contain the hundreds of tandemly repeated copies of ribosomal RNA genes needed to support cellular viability. This repetitiveness makes it highly susceptible to copy number (CN) loss, threatening multi-generational maintenance of rDNA. How this threat is counteracted to avoid extinction of the lineage has remained unclear. Here, we show that the rDNA-specific retrotransposon *R2* is essential for rDNA CN maintenance in the *Drosophila* male germline, despite the perceived disruptive nature of transposable elements. Depletion of *R2* led to defective rDNA CN maintenance, causing a decline in fecundity over generations and eventual extinction. This study reveals that active retrotransposons can provide a benefit to their hosts, contrary to their reputation as genomic parasites, which may contribute to their widespread success throughout taxa.

**One Sentence Summary:** The retrotransposon *R2* initiates restoration of ribosomal DNA copies to trans-generationally maintain essential locus.

## Main Text

Ribosomal RNAs (rRNAs) account for 80-90% of all transcripts in eukaryotic cells(*1*). To meet this demand, the ribosomal DNA (rDNA) gene that codes for rRNA is tandemly repeated hundreds of times, comprising rDNA loci on eukaryotic chromosomes. Ironically, this repetitive structure is susceptible to intra-chromatid recombination that causes rDNA copy number (CN) loss **(Fig. 1A)**, which is a major cause of replicative senescence in budding yeast (*2*). Evidence of similar rDNA CN instability has been noted in some tissues from aged dogs and humans(*3*, *4*). Critically, age-associated rDNA CN loss also occurs in the *Drosophila* male germline and is inherited by the next generation(*5*). The essential yet unstable nature of rDNA raises the question as to how the degeneration of rDNA loci over successive generations is prevented to avoid the extinction of the lineage. Intensive studies have revealed that sister chromatid recombination mediates rDNA CN expansion in yeast, thereby maintaining rDNA repeat abundance over generations(*2*). rDNA CN is variable between individuals of most species but maintained within a consistent range throughout the population(*6*), implying that transgenerational dynamic CN changes (loss and restoration) are a common feature of rDNA maintenance. However, the mechanism of transgenerational rDNA CN maintenance in multicellular organisms has remained a mystery. Over 50 years ago, the phenomenon of ‘rDNA magnification’ in *Drosophila* was first described as the process wherein aberrant rDNA loci bearing minimal rDNA repeats recover to a normal rDNA CN(*7*, *8*). rDNA magnification requires genes involved in homologous recombination-mediated repair, similar to yeast, which can duplicate tandemly repeated elements(*9*). Despite the robust rDNA CN expansion activity, it remained unclear if the mechanisms of rDNA magnification served a physiological function in natural populations. We recently demonstrated that offspring who inherited reduced rDNA CN from old fathers could also recover rDNA CN in their germline(*5*). This recovery depends on the same set of the genes as rDNA magnification, leading us to propose that rDNA magnification is a manifestation of the physiological mechanisms to maintain rDNA CN across generations(*6*). However, the underlying factors responsible for rDNA magnification remains poorly understood.

**Figure 1:**
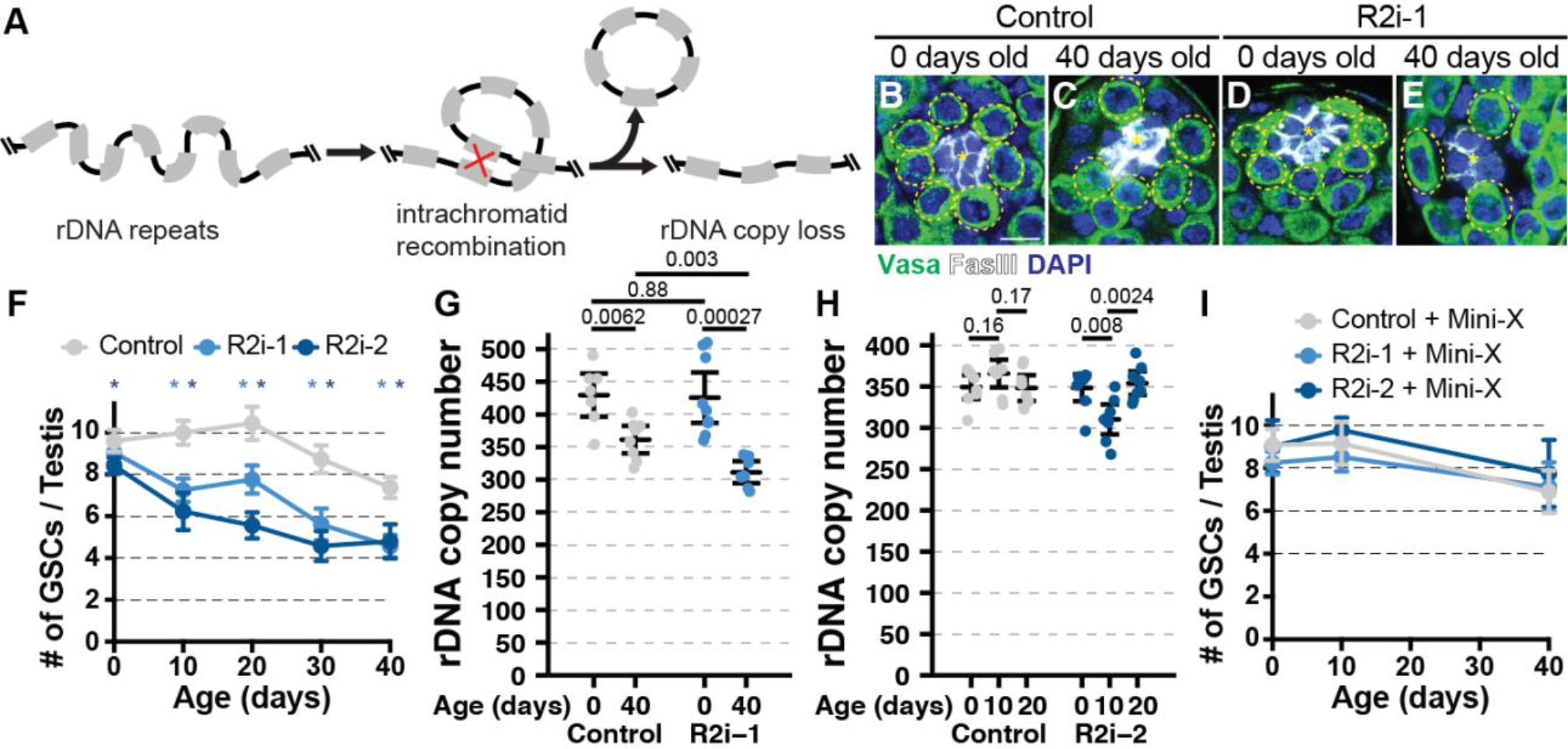
*R2* is required for GSC maintenance via rDNA CN maintenance during *Drosophila* male germline aging. **(A)** Model of rDNA repeat instability. **(B-E)** GSCs in 0- and 40-day old control (B-C) and *R2* RNAi testes (D-E). Yellow dotted circle = GSC. GSC signaling niche indicated by *. Green = Vasa, White = Fascillin III, Blue = DAPI. Scale bar = 7.5 μM **(F)** Average GSCs per testis in control and two *R2* RNAi lines during aging. * indicates p < 10^−3^ determined by Student’s t-test. Error = 95% confidence interval (CI). **(G-H)** Testis rDNA CN determined by ddPCR. P-value by Student’s t-test. Error = 95% CI. **(I)** Average GSCs per testis during aging in Control and two *R2* RNAi lines containing mini-X chromosome. Error = 95% CI.

Metazoan rDNA genes are frequently inserted by rDNA-specific transposable elements (TEs), such as the retrotransposon *R2* in *Drosophila*. *R2* is found throughout arthropods and *R2*-like elements are widely present across taxa, including Cnidaria, Planaria, nematodes, fish, birds, and reptiles(*10*, *11*). These TEs use their sequence-specific nuclease to mobilize specifically within rDNA loci(*12*), inserting into rDNA genes and likely disrupting 28S rRNA function(*13*) **(Fig. S1A)**. Surprisingly, we found that RNAi-mediated knockdown of *R2* in the *Drosophila* male germline (*nos-gal4*-driven expression of RNAi lines, *nos*>*R2i-1* or *R2i-2*, hereafter) **(Fig. S1A-C)** resulted in premature loss of germline stem cells (GSCs) during aging **(Fig. 1B-F)**. GSCs continuously produce differentiating germ cells to sustain sperm production throughout adulthood and thus are the source of the genome passed to the next generation(*14*). Whereas newly eclosed *R2* RNAi males contained similar numbers of GSCs to controls, GSC number more rapidly declined during aging in *R2* knockdown males compared to controls **(Fig. 1F)**. Given that *R2* is specifically inserted into rDNA, we also examined the effect of *R2* knockdown on rDNA stability.

Using highly quantitative droplet digital PCR (ddPCR), we found that RNAi-mediated knockdown of *R2* enhanced rDNA CN loss in the testis during aging **(Fig. 1G-H)**. Interestingly, one of the RNAi constructs (*R2i-2*) suffered rapid rDNA CN loss within the first 10 days of adulthood, but recovered by 20 days of age **(Fig. 1G)**. As GSC number was also initially more drastically affected with this RNAi line **(Fig. 1F)**, we speculate that severe rDNA loss caused by this RNAi construct (*R2i-2*) may select for GSCs that are less sensitive to *R2* RNAi activity. rDNA CN loss in the germline was further confirmed by DNA FISH on the meiotic chromosomes **(Fig S2A-F)**. rDNA CN insufficiency is likely the primary cause of GSC loss in *R2* RNAi animals, because increasing total rDNA CN via introduction of a mini-chromosome harboring an rDNA locus(*15*) suppressed the premature GSC loss caused by *R2* knockdown **(Fig. 1I)**. These results revealed that *R2* contributes to sustaining GSC population during aging through rDNA CN maintenance, uncovering an unanticipated benefit of the *R2* retrotransposon to the host, despite the widely-held view of TEs being genetic parasites.

Why is *R2* necessary for rDNA copy number maintenance? We found that *R2* is required for rDNA magnification. rDNA magnification is detected as the emergence of offspring with normal cuticle from fathers with abnormal (‘bobbed’) cuticle caused by insufficient rDNA CN(*7*) **(Fig. 2A)**. *Drosophila melanogaster* rDNA loci reside on the sex chromosomes (X and Y)(*16*), and rDNA magnification almost exclusively occurs to X chromosome rDNA loci harboring the minimal viable amount of rDNA (*bb^Z9^*, **Fig. S3A)** when combined with a Y chromosome lacking rDNA (*bb^Z9^/Ybb^0^*, ‘magnifying males’ hereafter)(*7*) **(Fig. S3B)**. Importantly, rDNA magnification never occurs in males with a normal Y chromosome containing intact rDNA (*bb^Z9^/Y^+^*, ‘non-magnifying males’ hereafter)(*8*), indicating the presence of mechanisms to monitor rDNA CN to activate expansion. We found that *R2* knockdown reduces rDNA magnification from 13.73% (control, n = 233) to 0% (*R2i-1*, n = 181, p = 5.6×10^−7^) and 2.36% (*R2i-2*, n = 127, p =9.9×10^−4^) **(Fig. 2B)**. Moreover, quantification of rDNA CN by ddPCR revealed that 87.5% of *bb^Z9^* chromosomes increased rDNA CN in magnifying males (n = 96, p = 1.8×10^−4^), with an average increase of 18.29 rDNA copies across all samples (n = 96, p = 3.1×10^−12^) **(Fig. 2C)**. This widespread rDNA CN increase indicates that rDNA magnification broadly increases rDNA CN throughout the germline, despite only 13.73% of *bb^Z9^* chromosomes recovering enough CN to support normal cuticle development. This rDNA CN increase in magnifying males is also eliminated upon *R2* knockdown **(Fig. 2C)**. These results reveal that *R2* is required for rDNA CN expansion during rDNA magnification. Interestingly, we found that rDNA magnification was blocked only when the *R2* RNAi constructs were expressed by the *nos*-*gal4* driver in early germ cells (including GSCs), but not when expressed in later germ cells by the *bam*-*gal4* driver (see Methods) **(Fig. 2B)**. These results indicate that *R2* primarily functions in the earliest stages of germ cells (including GSCs) to support rDNA magnification.

**Figure 2:**
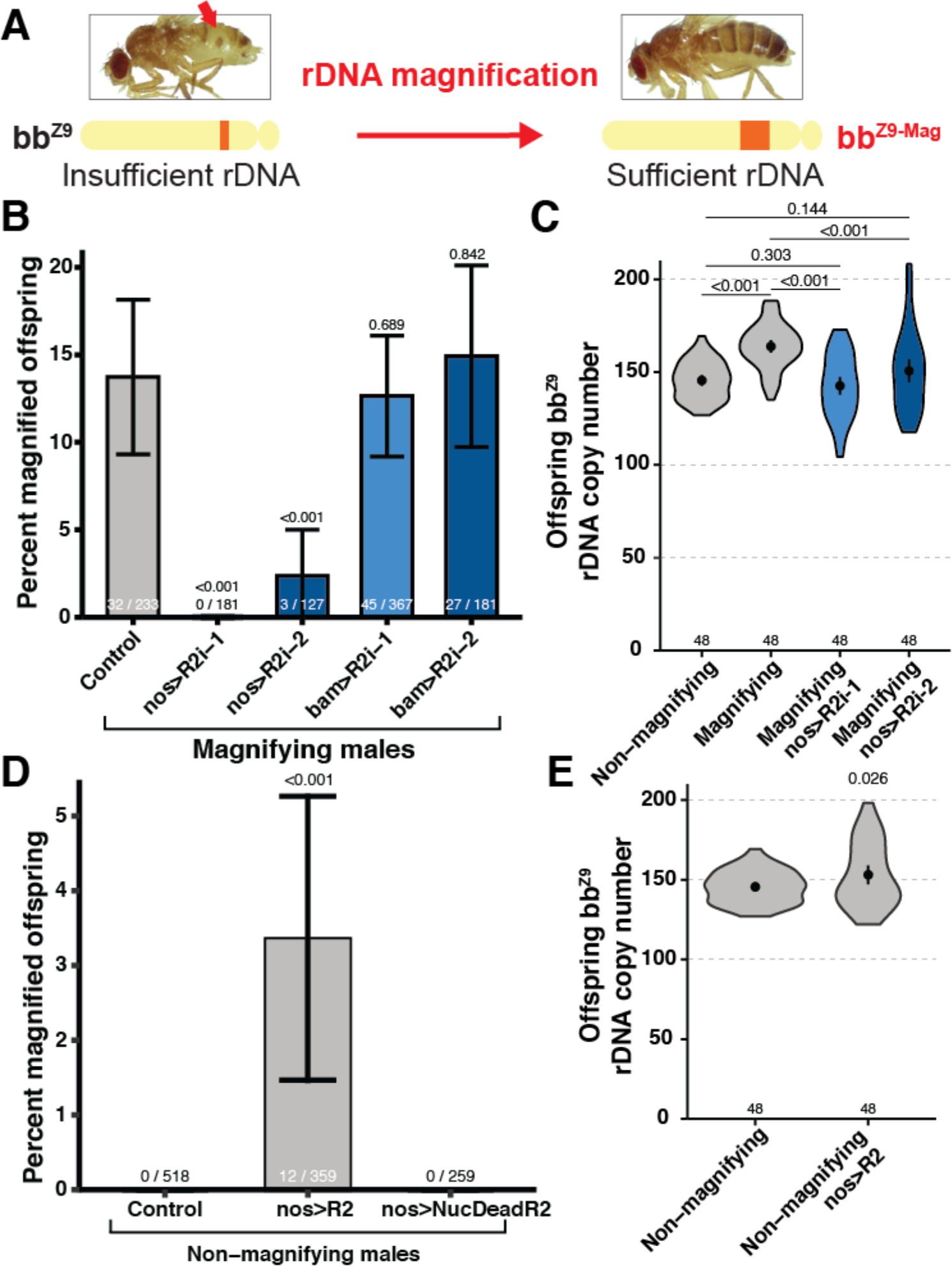
*R2* is necessary and sufficient for rDNA magnification. **(A)** Diagram of rDNA magnification at the *bb^Z9^* rDNA locus, during which dorsal cuticle defect (red arrow) revert to normal cuticle. **(B)** Percent magnified offspring determined by cuticular phenotype in offspring from magnifying males. P-value determined by chi-squared test. Error = 95% CI. **(C)** Mean *bb^Z9^* locus rDNA CN determined by ddPCR in daughters from males. P-value determined by Student’s t-test. Error = 95% CI. **(D)** Percent magnified offspring from non-magnifying males. P-value determined by chi-squared test. Error = 95% CI. **(E)** Mean *bb^Z9^* locus rDNA CN determined by ddPCR in daughters from non-magnifying males. Non-magnifying condition is the same data as panel (C). P-value determined by Student’s t-test. Error = 95% CI.

We further found that *R2* is sufficient for rDNA CN expansion. Ectopic expression of transgenic *R2* in the germline **(Fig. S4A-F)** induced rDNA magnification of the *bb^Z9^* locus in non-magnifying males (*bb^Z9^/Y^+^*) **(Fig. 2D-E)**. While rDNA magnification assessed by dorsal cuticle is never detected from control males (*bb^Z9^/Y^+^* without *R2* expression), we found 3.3% of male offspring exhibited magnification (normal cuticle) upon expression of transgenic *R2* **(Fig. 2D**, n = 877, p = 3.2×10^−5^). Importantly, reversion of the cuticle phenotype was heritable to the subsequent F2 generation throughout our experiments, confirming that CN restoration occurred in the germline **(Fig. S5A-C)**. Quantification of rDNA CN by ddPCR revealed that ectopic overexpression of *R2* in non-magnifying males (*bb^Z9^/Y^+^*) also increases the average rDNA CN at *bb^Z9^* rDNA loci among all offspring **(Fig. 2E**, n = 94, p = 0.0256), revealing *R2* is sufficient to induce rDNA CN expansion. Critically, expression of a nuclease dead *R2* transgene (NucDead*R2*) in non-magnifying males **(Fig. S4A-E)** failed to induce rDNA magnification **(Fig. 2D)**, suggesting that the nuclease activity of *R2* is essential for its ability to induce rDNA CN expansion.

How does *R2*’s nuclease activity contribute to rDNA magnification? In yeast, rDNA CN expansion is initiated by double-stranded breaks (DSBs) at the rDNA intergenic sequence, which induces sister chromatid recombination that results in rDNA gene duplication(*17*). All proposed models of *Drosophila* rDNA CN expansion (the most prominent model being unequal sister chromatid recombination(*18*)) require an initiating DSB at the rDNA locus, **(Fig. 3A; Fig. S6A-B)**. Indeed, artificial introduction of DSBs at rDNA loci by I-CreI endonuclease expression has been reported to induce rDNA magnification(*19*), but the endogenous factor that induces rDNA magnification remained unclear. *R2* is capable of creating DSBs through sequential nicking of opposite DNA strands during retrotransposition(*10*). It has been speculated that DSBs created during *R2* retrotranspostion may be an initiating event of rDNA magnification(*9*), although this possibility has yet to be empirically tested. We confirmed that ectopic overexpression of *R2*, but not NucDead*R2*, indeed induces chromosomal breaks at rDNA loci identified by chromosome spreads **(Fig. S4B-D)**. *R2* overexpression (but not NucDead*R2*) in the germline also increased the frequency of GSCs with DSBs, identified by γH2Av expression **(Fig. S4E)**. Next, we found rDNA magnification is associated with an elevation in DSBs in GSCs: the frequency of γH2Av-positive GSCs is increased in magnifying males (*bb^Z9^/Ybb^0^*) compared to non-magnifying males (*bb^Z9^/Y^+^*) (**Fig. 3B-C, E**; n = 519, p = 8.8×10^−4^). Strikingly, we observed that knockdown of *R2* in magnifying males reduced the frequency of γH2Av-positive GSCs to levels comparable to non-magnifying males **(Fig. 3D-E;** n = 537, p = 7.1×10^−4^ for *R2i-1*; n = 521, p = 7.9×10^−4^ for *R2i-2*), indicating that *R2* is responsible for the DSBs formed in GSCs during rDNA magnification. Taken together, these results suggest that rDNA-specific endonuclease activity of *R2* creates DSBs at the rDNA loci that may in turn induce rDNA CN expansion.

**Figure 3:**
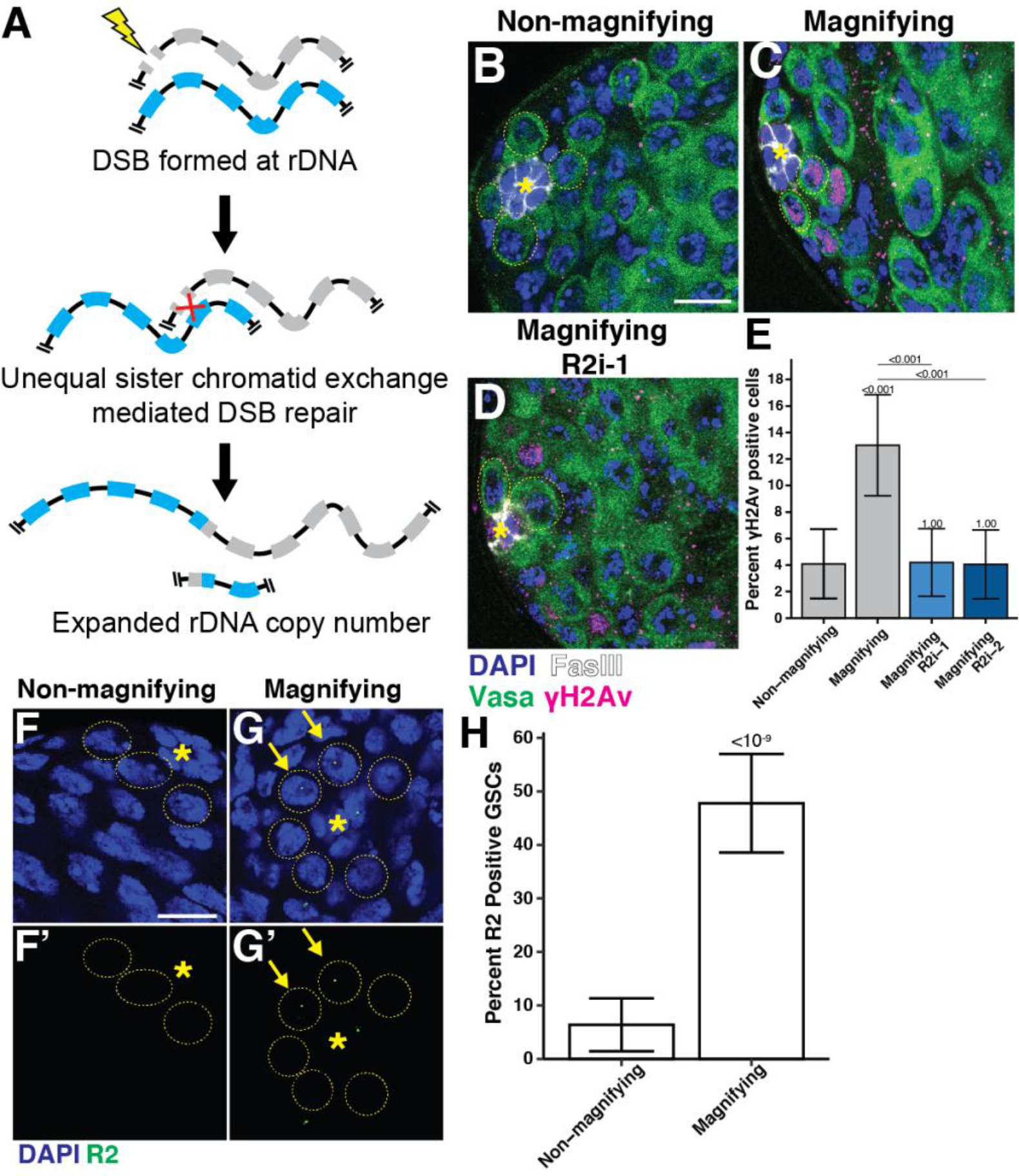
Derepressed *R2* creates DSBs in GSCs during rDNA magnification. **(A)** Diagram of rDNA CN expansion by unequal sister chromatid exchange during DSB repair at rDNA loci. Recombination between misaligned rDNA copies during DSB repair result in crossovers that create unequal sister chromatid exchange that increases rDNA CN on one chromatid. **(B-D)** Detection of DSBs in the early adult male germline by anti-γH2Av staining. *R2* RNAi expressed under the *nos-gal4* driver. Non-RNAi conditions contain the *nos-gal4* driver alone. GSCs indicated by yellow dotted circle. Blue = DAPI, Green = vasa, Magenta = γH2Av, white = FasIII. The hub is indicated by *. Scale bar = 10 μM. **(E)** Percentage of γH2Av positive GSCs. P-value determined by chi-squared test. Error = 95% CI. **(F-G)** *R2* expression in GSCs (yellow dotted circle). Blue = DAPI, Green = *R2* mRNA. Isolated *R2* channel in F’-G’. The hub is indicated by *. *R2* positive cells GSCs are marked by yellow arrowhead. **(H)** Percentage of *R2* positive GSCs. P-value determined by chi-squared test. Error = 95% CI.

Given the threat *R2* mobilization poses to the host genome, both by disruption of rRNA function and causing excessive DSB formation(*10*), how is the potential benefit of *R2* to rDNA CN maintenance balanced with the detriment of *R2* retrotransposition? We found *R2* expression in the germline is specifically de-repressed under conditions of reduced rDNA CN, potentially explaining how the conflicting consequences of *R2* expression are resolved. Using RNA fluorescence *in situ* hybridization (RNA FISH) to examine *R2* expression at a single cell resolution, we found that the frequency of GSCs expressing *R2* was significantly increased in magnifying males (*bb^Z9^/Ybb^0^*), whereas non-magnifying males (*bb^Z9^/Y^+^)* rarely expressed *R2* **(Fig. 3F-H**; n = 231, p = 1.7×10^−10^). Moreover, we found that GSCs from aged animals and the sons of old fathers, which inherit reduced rDNA CN(*5*),also exhibited a higher frequency of *R2* expression compared to GSCs from young flies **(Fig. S7A-B, D**; n = 1,247, p = 8.3×10^−4^ for old animals; n = 1,107, p = 1.5×10^−4^ for offspring). Importantly, the frequency of *R2* expression among GSCs in the sons of old fathers returned to the basal level after 20 days of age, when rDNA CN was shown to have recovered(*5*) **(Fig. S7C-D**; n = 617, p = 0.036). These results indicate that *R2* expression is dynamically regulated in response to changing rDNA CN. Taken together, we propose that *R2* expression is finely tuned to function when most beneficial to the host while minimizing unnecessary exposure to the harmful effects of transposition.

Based on the finding that *R2* plays a critical role in maintaining germline rDNA CN, we postulated that *R2* is essential to prevent continuous multi-generational rDNA loss capable of causing the extinction of the lineage. In *C. elegans*, the loss of genome integrity is known to cause gradual loss of fertility, a phenotype known as <u>mor</u>tal <u>g</u>ermline (morg)(*20*). To test whether *R2*-mediated rDNA maintenance is required to maintain fertility through generations, we established multiple independent lines expressing *R2* RNAi in their germline and tracked their fecundity at each generation through the ability of each line to produce sufficient offspring to establish a new generation **(Fig. S8)**. While nearly all control lines survived throughout the duration of the experiments, we found that lines expressing the *R2i-1* RNAi construct failed to consistently produce sufficient progeny, with over half failing by the fourth generation **(Fig. 4A)** (n = 43, p = 3×10^−6^), indicating that *R2* is essential for continuity of the germline lineage. Surviving males of extinguishing *R2i-1* lineages had ~20% reduction in rDNA CN compared to control lines (n = 22, p = 0.031) **(Fig. 4B)**. With the *R2i-2* RNAi, the lineage was maintained relatively well, after initial sharp drop **(Fig. 4A)**: Considering that *R2* knockdown by the *R2i-2* construct exhibits only transient rDNA CN decrease at day 10 (**Fig S2A**), we speculate that this RNAi construct quickly selects for the germ cells and lineages insensitive to *R2* knockdown. Taken together, these results suggest that *R2*-mediated maintenance of rDNA contributes to germline immortality.

**Figure 4:**
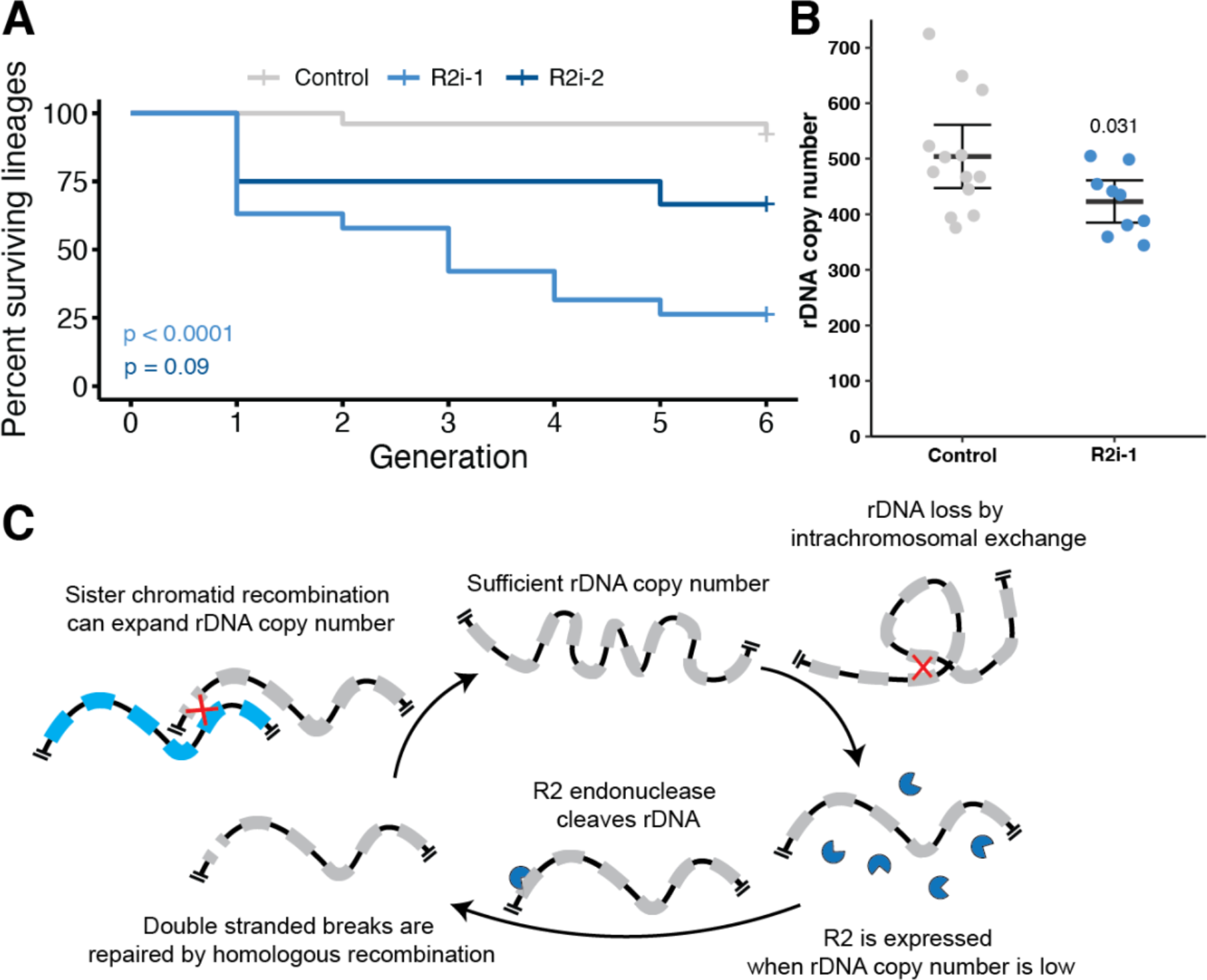
*R2* is required to maintain rDNA CN and fertility over successive generations. **(A)** Kaplan-Meier curve of lineage survival in control (*nos-gal4* driver alone and two *R2* RNAi expressing (via the *nos-gal4* driver) lineages. Each lineage constitutes an individual data point. P-values determined by log rank test. **(B)** rDNA CN determined by ddPCR in males of control animals at the 6^th^ generation or *R2* RNAi lineages at their terminating generation. P-value determined by Student’s t-test. Error = 95% CI. **(C)** Model of the role of *R2* in germline rDNA CN maintenance.

Together, our findings reveal an unanticipated ‘function’ of retrotransposon activity to benefit the host genome through a role in rDNA CN maintenance. Repetitive DNA sequences are among the most vulnerable elements of the eukaryotic genome(*21*). We propose that DSBs generated by *R2* in GSCs with reduced rDNA CN stimulate sister chromatid exchange that results in rDNA CN expansion **(Fig. 4C)**. Our proposed ‘function’ for *R2* in rDNA maintenance may represent a novel mutualistic host-TE relationship, which are rarely described in eukaryotes(*22*). *R2*’s de-repression when it can be beneficial (i.e. decreased rDNA CN) may be the key to this mutualistic host-TE relationship. It awaits future investigation to understand how the mechanisms that repress *R2*(*10*) may tune this activity for host’s benefit. The widespread presence of *R2* and other rDNA-specific TEs in both vertebrates and invertebrates(*11*) suggests that similar host-TE mutualism may support rDNA CN maintenance throughout Metazoa. Interestingly, many of rDNA-specific TEs have little sequence similarity to *R2*, instead appearing to be derived from other non-specific TEs(*11*), suggesting this host-TE mutualism may have evolved multiple times over the course of evolution. In summary, our study provides an example of mutualistic retrotransposons in the maintenance of eukaryotic genomes, and we propose that more mutualistic TEs are yet to be discovered.

## Acknowledgements

We thank the Bloomington *Drosophila* Stock Center, Kyoto *Drosophila* Stock Center and Developmental Studies Hybridoma Bank for reagents. We thank the Yamashita lab members and Dr. Andy Clark for discussion and comments on the manuscript. We thank ATCC for design of the 28S and RpL ddPCR assays.

## Authors contributions

JON: Conceptualization, Methodology, Verification, Formal analysis, Investigation, Resources, Data curation, Writing – Original draft, Writing – Review & editing, Visualization, Funding acquisition. AS: Validation, Investigation, Resources, Data curation, Writing – Review & editing. YMY Conceptualization, Methodology, Resources, Writing – Original draft, Writing – Review & editing, Visualization, Supervision, Project administration, Funding.

## Data materials availability

All data is available in the manuscript or the supplementary materials.

## List of Supplementary Materials

### Supplementary Materials

Materials and Methods

Fig S1 – S8

Table S1 – S2

References 23 - 30

