## Supplemental data and methods for "The retrotransposon *R2* maintains *Drosophila* ribosomal DNA repeats"

### Supplementary Materials

### Materials and Methods

*Immunofluorescence*

Immunofluorescence staining of testes was performed as previously described(*23*). Briefly, testes were dissected in PBS, fixed in 4% formaldehyde in PBS for 30 minutes, then briefly washed two times in PBS containing 0.1% Triton-X (PBS-T), followed by washing in PBS-T for 30 minutes. After washes, samples were incubated at 4°C overnight with primary antibody in 3% bovine serum albumin (BSA) in PBS-T. Samples were washed three consecutive times for 20 minutes in PBS-T, then incubated at 4°C overnight with secondary antibody in 3% BSA in PBS-T, washed three times again in PBS-T for 20 minutes, and mounted in VECTASHIELD with DAPI (Vector Labs). The following primary antibodies were used: rat anti-vasa (1:20; DSHB; developed by A. Spradling), mouse anti-Fascillin III (1:200; DSHB; developed by C. Goodman), and rabbit anti-γ-H2AvD pS137 (1:200; Rockland). Images were taken with a Leica Stellaris 8 confocal microscope with 63x oil-immersion objectives and processed using Fiji (ImageJ) software.

*RNA FISH*

RNA FISH samples were prepared as previously described(*5*). In short, dissected testes were fixed in 4% formaldehyde in PBS for 30 minutes, briefly washed in PBS, and permeabilized in 70% ethanol overnight at 4°. Samples were then briefly rinsed in 2xSSC with 10% formamide prior to hybridization with 50nM Stellaris probes overnight at 37°. Samples were washed twice in 2xSSC with 10% formamide for 30 minutes and mounted in VECTASHIELD with DAPI (Vector Labs). Samples were imaged using a Leica Stellaris 8 confocal microscope with 63x oil-immersion objectives and processed using Fiji (ImageJ) software.

*DNA isolation*

Testis DNA was isolated from 50 pooled dissected testes frozen in liquid N2. DNA isolation was performed according to previously described methods for isolation from Drosophila tissues(*24*). DNA was isolated from individual Drosophila animals using a modified protocol of the DNeasy Blood and Tissue DNA extraction kit (Qiagen). In short, individual animals were homogenized in 200 µL Buffer ATL containing proteinase K using a pipette tip in Eppendorf tubes, vortexed for 15 seconds, and incubated for 1.5 hours at 56º. Samples were then prepared following the manufacturer's protocol after incubation. All DNA samples were quantified and checked for purity by Nanodrop One spectrophotometer (ThermoFisher).

*rDNA copy number measurement by Droplet Digital PCR (ddPCR)*

30 ng of DNA sample were used per 20 µL ddPCR reaction for control gene reactions (RpL and Upf1), and 0.3 ng of DNA per 20 µL ddPCR reaction for 28S rDNA reactions. Primers and probes for reactions are listed at **Table S1**. ddPCR reactions were carried out according to the manufacturer's (Bio Rad) protocol. In short, master mixes containing ddPCR Supermix for Probes (No dUTP) (Bio Rad), DNA samples, primer / probe mixes, and HindIII-HF restriction enzyme (New England Biolabs) for 28S rDNA reactions (no restriction enzyme for control gene reactions) were prepared in 0.2 mL Eppendorf tubes, and incubated at room temperature for 15 minutes to allow for restriction enzyme digestion. ddPCR droplets were generated from samples using QX200 Droplet Generator (Bio Rad) and underwent complete PCR cycling on a C100 deep-well thermocycler (Bio Rad). Droplet fluorescence was read using the QX200 Droplet Reader (Bio Rad). Sample copy number was determined using Quantasoft software (Bio Rad). rDNA copy number per genome was determined by 28S sample copy number multiplied by 100 (due to the 100x dilution of sample in the 28S reaction compared to control reaction) divided by control gene copy number multiplied by the expected number of control gene copies per genome (2 for RpL in all samples; 2 for Upf1 in female samples; 1 for Upf1 in male samples). The 28S copy number

values determined by each control gene was averaged to determine 28S copy number for each sample.

*RNA isolation*

50 dissected testes were pooled and frozen in liquid N2 for each RNA isolation sample. Samples were homogenized in 400 µL TRIzol™ (ThermoFisher Scientific) and RNA was isolated using Direct-zol™ RNA Miniprep kit (Zymo Research) according to manufacturer directions, including on-column DNase I treatment. All RNA samples were quantified and checked for purity by Nanodrop One spectrophotometer (ThermoFisher).

*Quantification of R2 expression by reverse transcriptase (RT)-ddPCR*

Approximately 20 ng of total RNA were used per 20 µL RT-ddPCR reaction for R2 reactions, and 0.2 ng of total RNA were used per 20 µL control gene (Tubulin) reaction. Tubulin primers and probe are listed at **Table S1**, and R2 primers and probe mix were designed by Bio Rad (Assay ID: dCNS858096478). ddPCR droplets were generated from samples using QX200 Droplet Generator (Bio Rad) and underwent RT-PCR and end point PCR on a C100 deep-well thermocycler (Bio Rad). Droplet fluorescence was read using the QX200 Droplet Reader (Bio Rad). RNA quantitation was determined using Quantasoft software (Bio Rad), and R2 counts were normalized to Tubulin concentration for all samples. Normalized R2 expression values were then set relative to the average R2 expression value in control conditions.

*Mitotic and meiotic chromosome spread, DNA FISH, and quantification*

Mitotic chromosome spreads in neuroblast cells, DNA FISH, and imaging were all done as previously described(*25*). In short, brains were dissected from male third instar larvae in PBS and fixed in 25µ µL of acetid acid and 4% formaldehyde in PBS. Samples were applied to Superfrost plus slides and manually squashed under a coverslip, then immediately frozen in liquid N2. After freezing slides were removed from N2, coverslip removed, and slides were dehydrated in 100% ethanol and dried at room temperature. DNA FISH hybridization was performed in 20 µL of 50% formamide, 10% dextran sulfate, 2X SSC buffer and 0.5 µM each probe applied directly to the sample on the slide and covered with a cover slip. Samples were incubated at 95° for 5 minutes, cooled and wrapped in parafilm, then incubated overnight at room temperature in a dark humid chamber. Coverslips were removed and slides were washed three times for 15 minutes in 0.1x SSC, dried, and then mounted in VECTASHIELD with DAPI (Vector Labs). Samples were imaged using a Leica Stellaris 8 confocal microscope with 63x oil-immersion objectives and processed using Fiji (ImageJ) software. Meiotic chromosome spreads were prepared from dissected testes and imaged in the same manner. Relative Y:X rDNA fluorescence quantification was determined as previously described (*5*). Probes used for this study are as follows: 359, 5'-AGGATTTAGGGAAATTAATTTTTGGATCAATTTTCGCATTTTTTGTAAG-3'-Cy5; $(TAGA)_6$-Cy5; IGS, 5'-AGTGAAAAATGTTGAAATATTCCCATATTCTCTAAGTATTATAGAGAAAAGCCATTTTAGTGAATGGA-3'-Alexa488; $(AATAC)_6$-Cy3; and $(AATAAAC)_6$-Cy3.

*Generation of rDNA deletion animals*

rDNA copy number loss was induced during larval development in *yw* / Y ;; *HS-I-CreI*, *Sb* / TM6B males with a *y, w* X chromosome by I-CreI expression as previously described(*19*). In brief, parental animals mated and laid eggs for three days, then removed from food. After one additional day of larval development, animals were exposed to 37° C heat shock for 45 minutes on two consecutive days. identify X chromosomes with significant rDNA copy number reduction (bb), adult males that experienced I-CreI expression were mated to $bb^{158}$ / *FM6* females, and virgin non-*FM6* daughters (bb / $bb^{158}$) were screened for the *bobbed* phenotype. 28 out of 946 non-*FM6* daughters screened were *bobbed*. To isolate potentially reduced rDNA loci and remove HS-I-CreI from the background, TM6B containing *bobbed* females were individually mated to wild-type males. Male offspring from each individual female candidates were subsequently individually mated to *bb*$^{158}$ / *FM6* females. Any mating that failed to produce non-FM6 daughters were eliminated (due to having the bb$^{158}$ and not the candidate bb chromosome). All viable non-FM6 daughters were double checked for the *bobbed* phenotype, and stocks with all *bobbed* non-FM6 daughters had FM6 containing siblings collected and used to establish a bb / FM6 stock. This method isolated the novel rDNA deletion allele, bb$^{Z9}$, used in this study.

*Drosophila genetics*

All Drosophila lines used in this study are found in **Table S2**. All animals we reared on standard Bloomington medium at 25º. All aging was done in roughly 1:1 mixed presence of males and females, provided fresh food every four to six days. *UAS-R2 RNAi* strains were designed using SNAP-DRAGON shRNA target software, and oligos containing target hairpin sequence were cloned into the WALLIUM20 vector for phiC31 site-directed integration into the Drosophila genome. The target sequence for the R2i-1 construct is 1481-CCGGTTGAACTCATCAATCAA-1502. The target sequence for the R2i-2 construct is 432- CCAGACGAACTTGATGAAGAA-453. All UAS-R2 transgenes were synthesized into pUAST:attB by VetcorBuilder (Chicago, IL) for site-directed integration. Importantly, these target sequences were specifically designed to target the R2 ORF encoding the R2 retrotransposase. ORF-containing R2 mRNA are expected to be translated in the cytoplasm, where it is subjected to silencing by the canonical RNAi mechanism. The UAS-R2 transgene contains the R2 ORF tagged with 3xFLAG tag at N-terminus, cloned into the pUAST:attB vector. The UAS-R2 transgene also contains sense mutations at the R2i-1 target sequence to render the transgene insensitive to this RNAi (1481-CCGGTTGAACTCATCAATCAA-1502 to 1481-ACGTCTTAATAGCAGTATTAA-1502. The Nuclease-Dead UAS-R2 transgene is identical to the UAS-R2 transgene except for 3001-AAACCAGAC-3009 to GCC and 3097-AAAATCAATAGA-3108 to 3097-GCCATCAATGCC-3108, which are analogous mutations to those demonstrated to disrupt *B. mori* R2 endonucelase activity(*26*). All injections and selection of animals containing integrated transgenes were performed by BestGene, Inc (Chino Hills, CA).

*rDNA magnification and heritability assays*

Males containing the $bb^{Z9}$ allele were mated in bulk to $bb^{158}$ / *FM6*, *Bar* females. $bb^{Z9}$ / $bb^{158}$ female offspring were selected based on the absence of the *Bar* dominant marker, and scored for cuticular phenotype. To determine the frequency of heritability of magnified offspring, unmated $bb^{Z9}$ / $bb^{158}$ female F1 animals with wild-type cuticles were collected and individually mated with 3 $bb^{158}$ / Y males. Since homozygous $bb^{158}$ animals are lethal, all viable female F2 animals are $bb^{Z9}$ / $bb^{158}$, and were scored for cuticular phenotype. Each individual F1 animal was scored for their ability to produce any offspring with wild-type cuticles, and for the percentage of their F2 female offspring to have wild-type cuticles.

*Lineage survival assay*

Independent lineages of *nos-gal4* / *CyO*; *UAS-R2 RNAi* / *Tm6B* or *nos-gal4* / *CyO*; *TM2* / *TM6B* animals were established by collecting siblings of the indicated genotypes from *nos-Gal4* / *CyO*; *TM2* / *TM6B* males mated to *Sp* / *CyO*; *UAS-R2 RNAi* / *TM6B* females. At each generation in each lineage, 3 males were mated with 5 females for 5 days, and offspring were collected 10 days after mated ended. Any lineages that did not have at least 3 males and 5 females at collection time were terminated due to insufficient animals.

*Statistics*

For all comparisons of percentage of samples with categorical values (percent γH2Av or R2 positive cells), significance was determined by chi-squared test. For all comparisons of samples with independent values (number of GSCs; rDNA copy number), significance was determined by Student's t-test between experimental and control conditions, unless otherwise indicated.

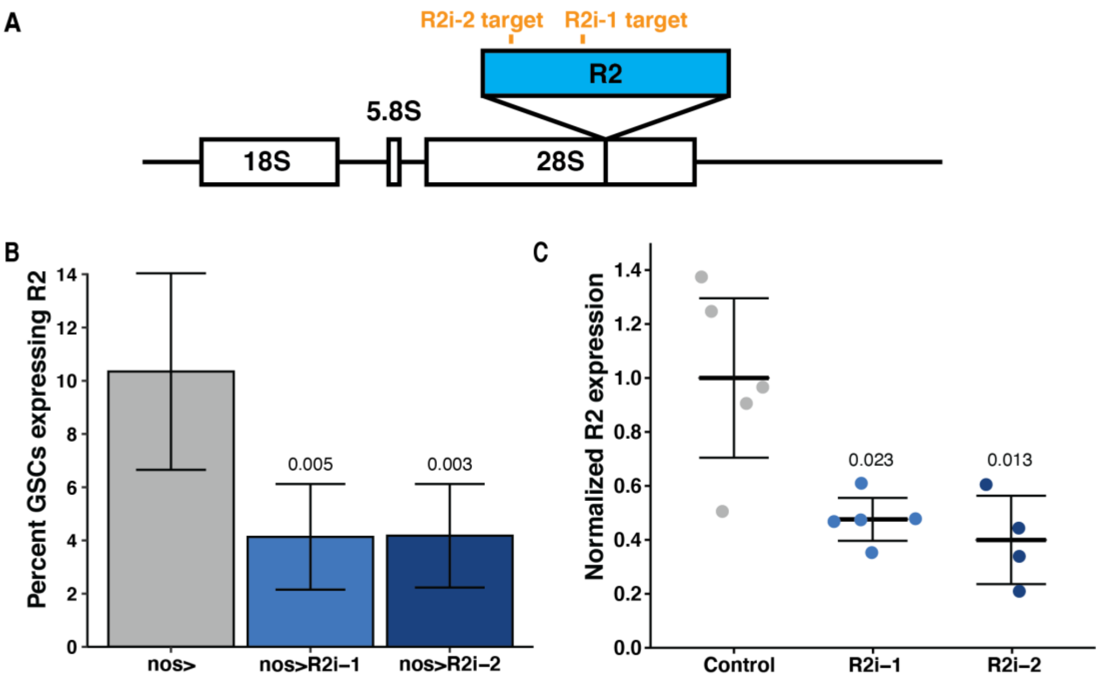


**Figure S1: Design and efficiency of R2 RNAi lines.** **(A)** Diagram the 45S rRNA cistron constituting an 'rDNA copy' and the sequence-specific R2 insertion site in the 28S rRNA gene. Location of R2 RNAi target sequences indicated in orange. **(B)** Precent of GSCs expressing R2 detected by RNA FISH in *nos-gal4* driver only condition (*nos*>) and *nos-gal4*-driven R2 RNAi (*nos>UAS-R2i*). Two independent R2 RNAi constructs were used throughout the study. P-value determined by chi-squared test. Error bars = 95% CI. **(C)** Efficiency of R2 knockdown used in this study determined by RT-ddPCR, normalized to Tubulin and set relative to control expression. GSCs were enriched by co-expressing *upd* (*27*, *28*), comparing *nos>upd* vs. *nos>upd, R2i*. P-value determined by Student's t-test. Error bars = 95% CI.

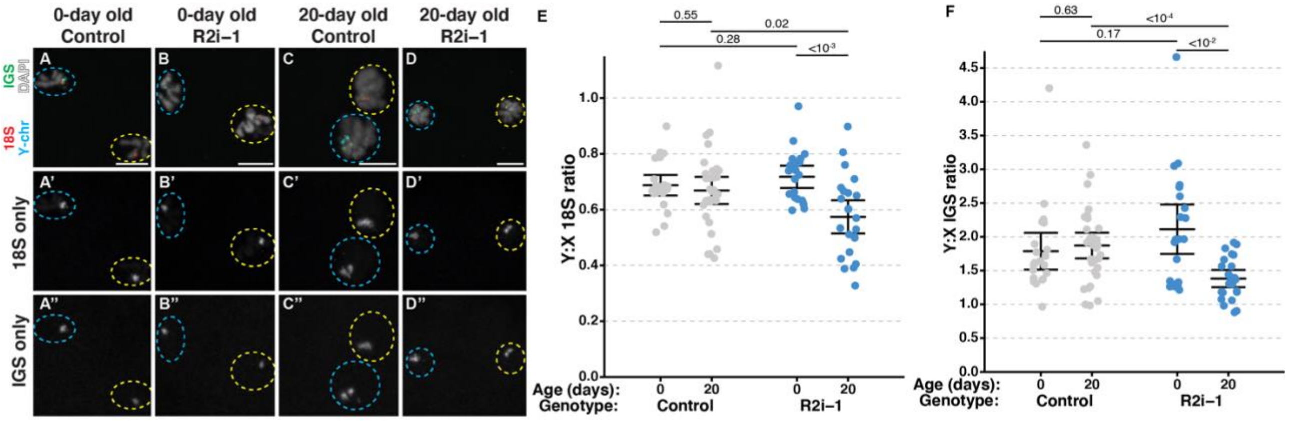


**Figure S2: Germline R2 inhibition causes rDNA CN loss in germ cells during aging. (A-D)** DNA FISH with 18S rDNA (Red), intergenic spacer (IGS) (Green) and the Y-specific AATAAAC (Cyan) probes on meiotic chromosome spreads from spermatocytes. Germline rDNA copy number loss during aging is biased on the Y chromosome, and thus results in a reduction in the relative Y:X ratio in rDNA (18S and IGS) FISH signal intensity (*5*). DAPI is shown in white. Y-containing chromatids are indicated by cyan dotted circle, X-containing chromatids are indicated by yellow dotted circle. A'-D': 18S (red channel), and A''-D'' IGS (green channel). Scale bar = 5 μm. **(E)** Relative Y:X 18S rDNA ratio of DNA FISH signal intensity. P-value determined by Student's t-test. Error = 95% CI from the mean. **(F)** Relative Y:X IGS ratio of DNA FISH signal intensity. P-value determined by Student's t-test. Error = 95% CI from the mean.

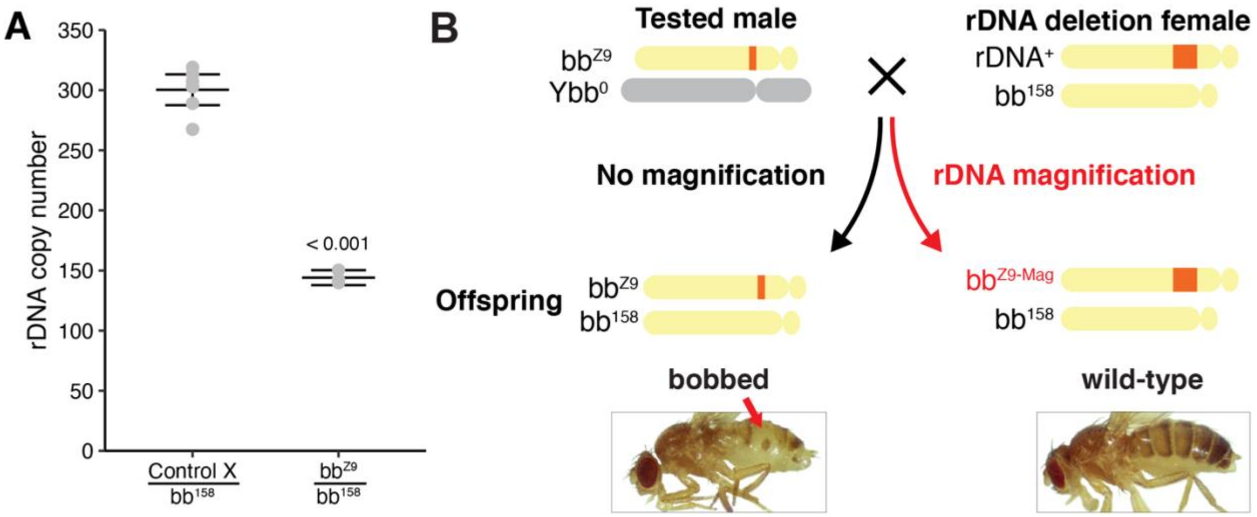


**Figure S3: Diagram of phenotypic assessment to detect rDNA magnification at the $bb^{Z9}$ allele. (A)** rDNA CN quantification of the $bb^{Z9}$ allele determined by ddPCR. All samples from individual females with either the $bb^{Z9}$ allele or X chromosome from which the allele was isolated heterozygous with an X chromosome completely lacking rDNA ($bb^{158}$). P-value determined by Student's t-test. Error bars = 95% CI. **(B)** Scheme to detect rDNA magnification of $bb^{Z9}$ allele. Tested magnifying males ($bb^{Z9}$ / $Ybb^{0}$) are crossed to females heterozygous for the $bb^{158}$ rDNA deletion allele. Resultant $bb^{Z9}$ / $bb^{158}$ daughters completely rely on the $bb^{Z9}$ locus for their rDNA. (Left) $bb^{Z9}$ / $bb^{158}$ offspring with the 'bobbed' cuticular defects associated with rDNA insufficiency (red arrow) are determined to have inherited a $bb^{Z9}$ allele that did not undergo rDNA magnification. (Right) $bb^{Z9}$ / $bb^{158}$ offspring with wild-type cuticles are determined to have inherited a $bb^{Z9}$ allele that restored normal rDNA content through rDNA magnification ($bb^{Z9\text{-}Mag}$). Thus, the portion of $bb^{Z9}$ / $bb^{158}$ offspring with wild-type cuticles represents the portion of $bb^{Z9}$ alleles that underwent rDNA magnification. $bb^{Z9}$ / $bb^{158}$ animals are also collected for rDNA CN quantification, and total rDNA CN in these animals represents the rDNA from the $bb^{Z9}$ locus alone.

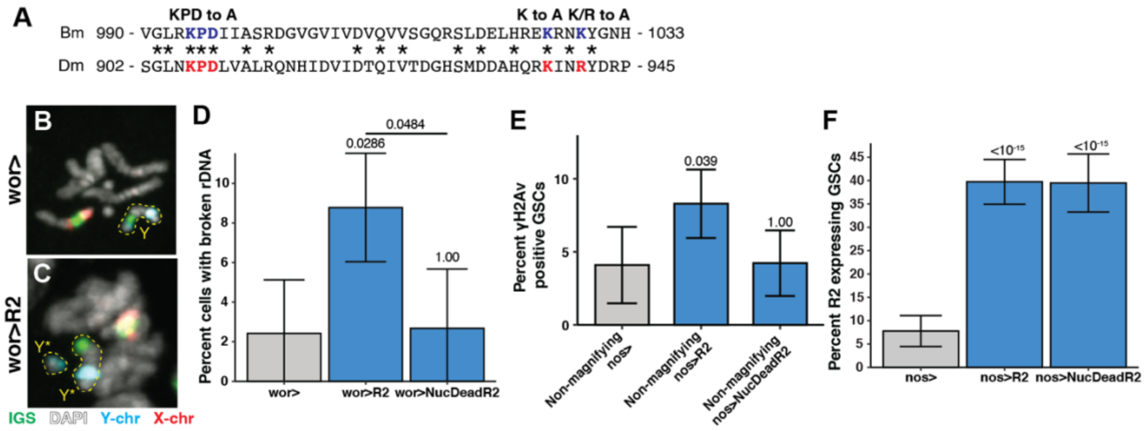


**Figure S4: Transgenic R2 expression induces double stranded breaks at rDNA loci. (A)** Alignment of 44 amino acids in the endonuclease domain of *Bombyx mori* R2 (Bm) and *Drosophila melanogaster* R2 (Dm). Residues shown to be essential for Bm R2 endonuclease activity(*26*) labeled in blue and respective Dm R2 residues in red. Mutations found to render Bm R2 endonuclease dead(*26*) and their respective amino acid changes in Dm R2 are indicated above highlighted residues. **(B-C)** DNA FISH in mitotic chromosome spreads of larval neuroblasts in animals with the neuroblast driver *wor-gal4* (wor>, B) or transgenic R2 expressed by the *wor-gal4* driver (wor>R2, C) animals. rDNA is labelled by probes targeting the rDNA intergenic sequence (IGS) (Green), Y- chromosome is labelled by probes targeting AATAC and AATAAAC satellite sequences (Cyan) and X-chromosome is labelled by probes against 359 and TAGA satellite sequences (Red). Y* indicates Y-chromosome fragments broken at the rDNA locus. **(D)** Percentage of neuroblasts with DSBs at rDNA loci determined by DNA FISH in mitotic spreads in indicated genotypes. P-values determined by chi-squared test. Error = 95% CI. **(E)** Percentage of γH2Av positive GSCs in the indicated genotypes. Transgenic R2 constructs expressed in the germline by the *nos-gal4* driver (*nos*>). P-value determined by chi-squared test. Error = 95% CI. **(F)** Percentage of GSCs expressing R2 determined by RNA FISH in control and R2 over-expression conditions. P-values determined by chi-squared test. Error = 95% CI.

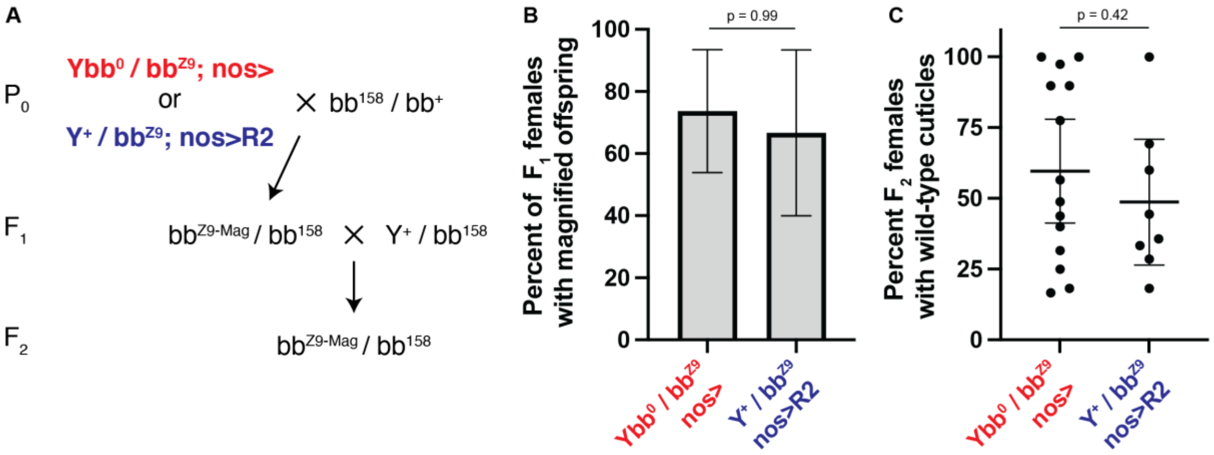


**Figure S5: rDNA magnification induced by insufficient rDNA copy number (*Ybb/bb*$^{z9}$) or ectopic R2 expression is stably inherited. (A)** Scheme to assess stable inheritance of rDNA magnification. $P_0$ Males capable of rDNA magnification (*Ybb*$^{0}$ */ bb*$^{Z9}$*; nos> and Y*$^{+}$ */ bb*$^{Z9}$; *nos>R2*) were mated to females heterozygous for a complete X-chromosome rDNA deletion (bb$^{158}$). Resultant putatively magnified *bb*$^{Z9\text{-}Mag}$ */ bb*$^{158}$ $F_1$ daughters with wild-type cuticles were mated with *bb*$^{158}$*/Y*$^{+}$ males, and the $F_2$ daughters *bb*$^{Z9\text{-}Mag}$ */bb*$^{158}$ were examined for their cuticle phenotype. $F_1$ animals that produced *bb*$^{Z9\text{-}Mag}$ */ bb*$^{158}$ $F_2$ daughters with wild-type cuticles were determined to have stably inherited rDNA magnification. **(B)** Frequency of $F_1$ females from indicated $P_0$ males that produced any $F_2$ animals with wild-type cuticles. Error = 95% CI. P-value by Chi-squared test. **(C)** Frequency of $F_2$ *bb*$^{Z9\text{-}Mag}$ */ bb*$^{158}$ offspring that have wild-type cuticles from stably magnified $F_1$ daughters from $P_0$ males of the indicated genotypes. The incomplete penetrance of wild-type cuticles among heritable rDNA magnification suggests epigenetic factors can influence the cuticular phenotype in animals with sub-optimal rDNA CN. Error = 95% CI. P-value determined by Student's t-test.

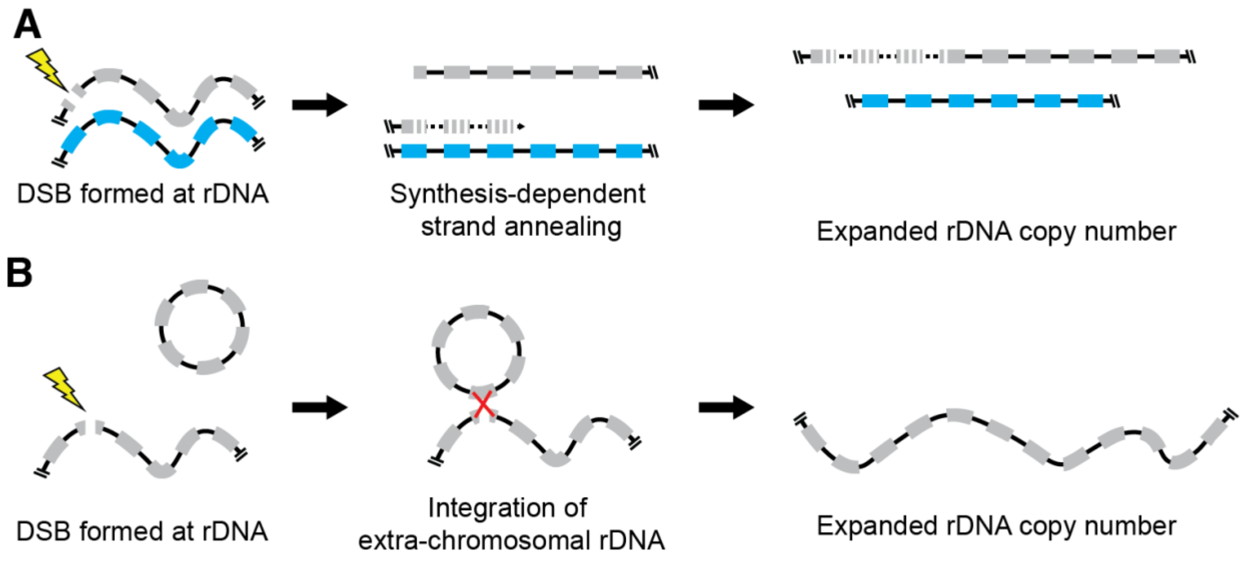


**Figure S6: Models of rDNA CN expansion initiated by DSBs at rDNA locus. (A)** rDNA CN expansion by strand-dependent strand annealing (SDSA). DSBs at rDNA loci can be repaired via recombination of a single strand with sister chromatid and synthesis. Synthesis can run on to replicate multiple rDNA copies before re-annealing with broken strand. Upon repair after synthesis, rDNA CN is expanded on one sister chromatid while CN of the other sister remains intact. **(B)** DSBs formed at rDNA loci induce homology-dependent repair that causes recombination with existing extrachromosomal rDNA circles (ERCs), allowing for reintegration of the ERC rDNA copies into the genomic rDNA locus.

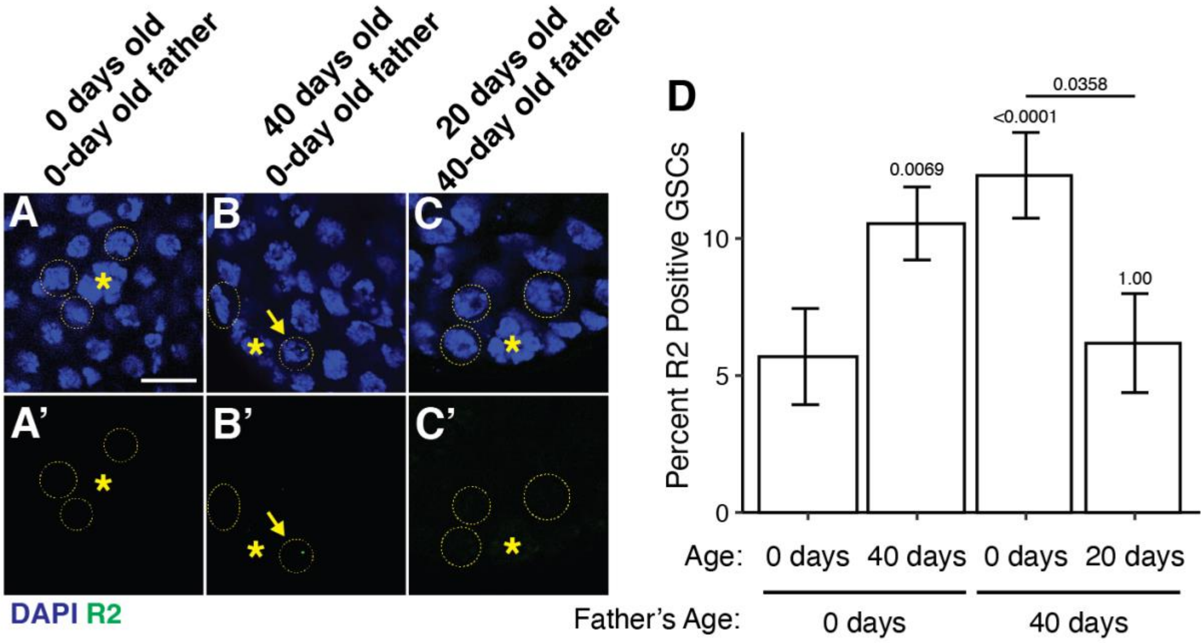


**Figure S7: R2 expression in GSCs is regulated in response to changing rDNA copy number during aging. (A-C)** R2 RNA FISH in the male germline of young (0-days old), (A) old (40-days old) (B), and recovered offspring of old animals (C). Isolated R2 channel in A'-C'. GSCs indicated by yellow dotted circle. * indicates GSC niche signal hub. R2 positive cells indicated by yellow arrowhead. **(D)** Percentage of R2 positive GSCs at indicated ages in the offspring of 0- or 40-day old fathers. P-value determined by chi-squared test. Error = 95% CI.

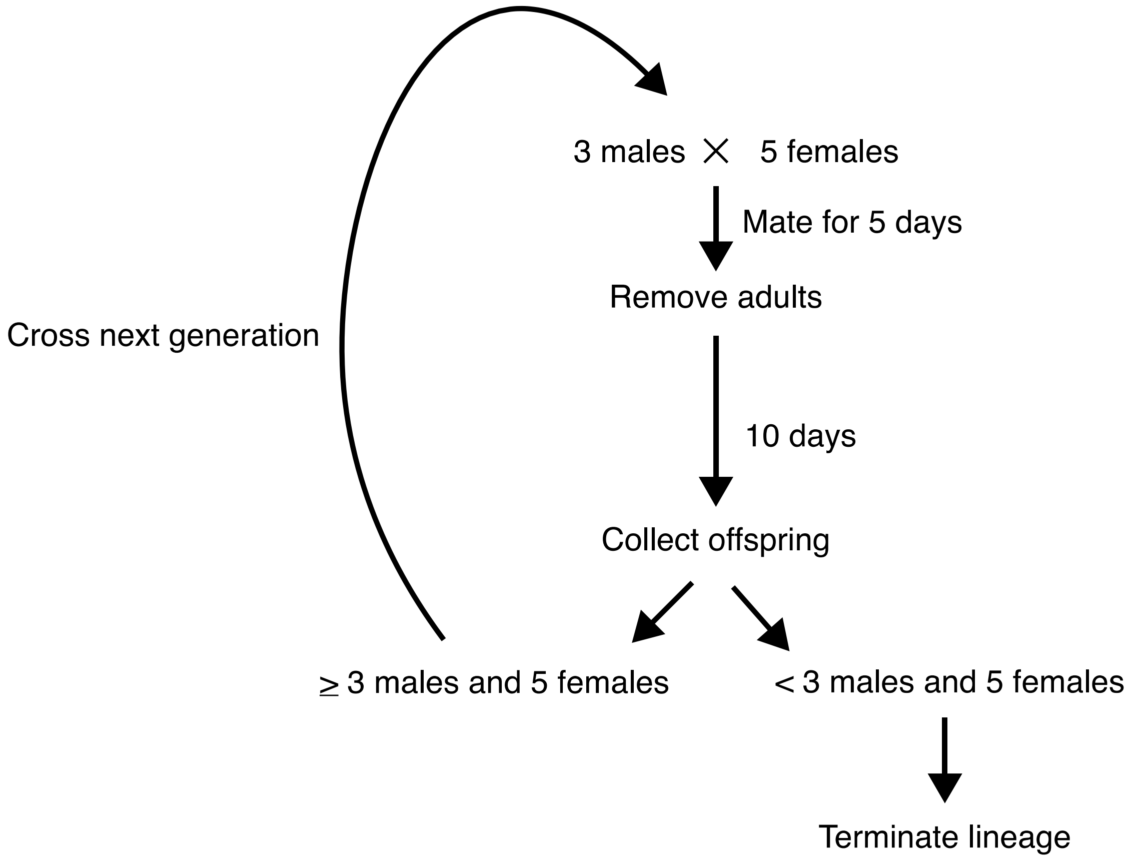


**Figure S8: Scheme for testing lineage survival in multi-generational R2 RNAi experiment.** Each lineage is established using 3 males and 5 females of the given genotype. Males and females are mated for 5 days and removed from the vial. 10 days later, all offspring are collected. If total offspring from the mating contains at least 3 males and 5 females, then 3 males and 5 females are randomly selected to establish the next generation of the lineage. If the total offspring contained fewer than 3 males or 5 females, then the lineage was terminated and future generations were not established. This procedure was repeated at each generation for each lineage for 6 generations or until termination.

**Table S1: List of oligomers used in this study for ddPCR assays.**

| Oligo Name | Type | Target | Sequence | Modifications |
|---|---|---|---|---|
| dd-RpL32 F | Primer | RpL32 | GCTTCAAGGGACAGTATCTG | |
| dd-RpL32 R | Primer | RpL32 | AACGCGGTTCTGCATGAG | |
| dd-RpL32 Probe | Probe | RpL32 | ATGCCCAACATCGGTTAC | 5' HEX AND 3' Iowa Black FQ |
| dd-28S F | Primer | 28S | GAGCTGCCATTGGTACAG | |
| dd-28S R | Primer | 28S | GCTTTCGCCTTGAACTTAG | |
| dd-28S Probe | Probe | 28S | TGGTGGATAGTAGCAAATAATCG | 5' 6-FAM AND 3' Iowa Black FQ |
| dd-Upf1 F | Primer | Upf1 | CACACTTTATGTCCACCATTATTG | |
| dd-Upf1 R | Primer | Upf1 | GAGTTTCCGTAGGGACCAC | |
| dd-Upf1 Probe | Probe | Upf1 | CCGTAACCGCCACTGCGGT | 5' 6-FAM AND 3' Iowa Black FQ |
| dd-Tubulin F | Primer | $\alpha$Tub84B | GAGCAGCTGATCACTGGTAAGG | |
| dd-Tubulin R | Primer | $\alpha$Tub84B | CGAGTGGAAGATGAGGAAGCCC | |
| dd-Tubulin Probe | Probe | $\alpha$Tub84B | CTGTCCAGAACCAGATCGACGATCTCCTTG | 5' 6-FAM AND 3' Iowa Black FQ |

**Table S2: List of Drosophila stocks used in this study.**

| Genotype | Source | Use |
|---|---|---|
| *Dp(1;f)122; C(1)RM, $y^1$ $v^1$ $f^1$ / C(1;Y)6, Df(1)259, $w^1$* | Kyoto Drosophila Genomics and Genetic Resource Center (DGRC) #107274 | Mini-x Chromosome (*Dp(1;f)*) |
| *yw* | Bloomington Stock Center (BSC) #1495 | Source for isolating novel rDNA deletion strain |
| *w[1118]; P{v[+t1.8]=hs-I-CreI.R}1A Sb[1]/TM6* | BSC #6937 | I-CreI endonucelase |
| *nos-gal4* | PMID: 9501989 | Early germ cell driver (*29*) |
| *wor-gal4* | BSC #56553 | Neuroblast driver |
| *bam-gal4* | PMID: 12571107 | Germ cell driver (*30*) |
| *y[1] eq[1]/Df(YS)bb[-]* | DGRC #101260 | Y chromosome complete rDNA deletion |
| *Df(1)bb158, y[1] / Dp(1;Y)y[+] / C(1)*; ca[1] awd[K]* | DGRC #106876 | X chromosome complete rDNA deletion |
| *UAS-upd* | PMID: 10346822 | Upd over expression to create GSC enrichment in testis |
